# Theanine boosts frontal theta and hippocampal beta and gamma oscillations for familiarity in object recognition

**DOI:** 10.1101/2025.05.08.652893

**Authors:** Kisa Watanabe, Kinjiro Takeda, Takeshi Nagahiro, Sena Iijima, Yuji Ikegaya, Nobuyoshi Matsumoto

## Abstract

Theanine, a non-proteinogenic amino acid derivative from green tea leaves, enhances object recognition memory in rats through facilitated neurogenesis. In this sense, the cellular mechanism for the theanine-enhanced object recognition memory has been elucidated to some extent, but physiological evidence still remains unclear. To tackle this issue, we chronically fed mice with theanine (or tap water) for three weeks, implanted electrodes into the hippocampus and frontal cortex, both of which are responsible for object recognition memory. We then recorded the local field potentials from the two regions during the novel object recognition task, evaluated the task performance, and broke down the neural signals in the hippocampus and frontal cortex into delta, theta, beta, low gamma, and high gamma frequency bands. The discrimination ratio of theanine-treated mice was higher than that of vehicle-treated mice. We also found that theta oscillations in the frontal cortex and beta and low gamma oscillations in the hippocampus in theanine-treated mice were simultaneously enhanced for familiar objects. These results shed light on the new physiological underpinnings of object recognition memory enhanced by exogenous substances.

## 1. Introduction

Object recognition memory is a subtype of episodic memory that promotes the ability to distinguish between new objects and previously encountered objects, thereby driving animals to update their perception of the external environment and act accordingly [1]. Earlier studies have demonstrated that lesion or inactivation of the hippocampus and perirhinal cortex impairs recognition memory [2,3], suggesting that neural pathways including these two regions supports object recognition memory in rodents [4,5]. From the theoretical point of view, the hippocampus and perirhinal cortex are believed to mediate processing of ‘where’ and ‘what’ information, respectively [6]. With this regard, it is widely accepted that the hippocampus and perirhinal cortex contribute to discrimination between objects.

Although there should not be direct connections between the hippocampus and perirhinal cortex, the two regions are connected with the medial prefrontal cortex that serves as a critical region for memory consolidation and decision-making [7]. In line with these studies, the medial prefrontal cortex is also involved in recognition memory [8,9]. Moreover, a recent study has determined that the hippocampus is more responsible for object recognition memory in mice than the perirhinal cortex when the mice sample identical objects in the training session for more than a few minutes [10].

While object recognition memory is affected by endogenous and exogenous substances that could modulate neural activity in the hippocampus and prefrontal cortex [11–15], few studies have specifically identified how these substances alter hippocampal and prefrontal neural activity for recognition memory based on electrophysiological parameters. For example, local field potentials (LFPs) recorded by extracellular electrodes reflect collective behavior of electric activity of individual neurons [16]. LFPs are often appraised in the form of oscillations in specific frequency bands such as delta, theta, alpha, beta, and gamma, each of which is functionally associated with vigilance states, locomotion, and cognition and affected by endogenous and exogenous substances [17–27].

Theanine (γ-glutamylethylamide), a unique amino acid component of green tea leaves, is a natural glutamate analog. After oral administration of theanine, it is absorbed from the gut, circulated in the body, and transported into the brain through the blood-brain barrier [28]. Due to its structural similarity to glutamate (and glutamic acid), theanine binds to and competitively antagonizes glutamate receptors [29–31]. In spite of relatively low affinity of theanine in binding to glutamate receptors [32], the glutamate receptor blockade by theanine regulates extracellular concentration of glutamate and counteracts excitotoxicity, which may lead to neuroprotective effects [33]; note that the neuroprotective effects of theanine can also be executed by suppression of mitochondrial radical formation although theanine does not function as a radical scavenger. Besides, theanine increases concentration of dopamine and γ-aminobutyric acid generally known as GABA [31,34] and modulates the noradrenaline and serotonin levels in the brain [35,36]. These regulatory effects of theanine on the neurotransmitter and neuromodulators has brought about the hypothesis that theanine modulates object recognition memory, which is demonstrated in rats; theanine facilitates object recognition memory [37–39]. However, it is poorly understood whether and how theanine coordinates neural activity in the hippocampus and frontal cortex underlying improvement of object recognition memory in mice.

To address this question, we recorded LFPs in the hippocampus and frontal cortex in mice that chronically treated with theanine and vehicle while they performed a novel object recognition task [13]. We then decomposed the LFPs into delta, theta, beta, and low and high gamma oscillations and assessed relevance of neural activity to memory performance.

## 2. Materials and methods

### 2.1 Ethical approval

Animal experiments were performed with the approval of the Animal Experiment Ethics Committee at the University of Tokyo (approval numbers: P4-6, P4-7) and according to the University of Tokyo guidelines for the care and use of laboratory animals. These experimental protocols were carried out in accordance with the Fundamental Guidelines for the Proper Conduct of Animal Experiments and Related Activities in Academic Research Institutions (Ministry of Education, Culture, Sports, Science and Technology, Notice No. 71 of 2006), the Standards for Breeding and Housing of and Pain Alleviation for Experimental Animals (Ministry of the Environment, Notice No. 88 of 2006) and the Guidelines on the Method of Animal Disposal (Prime Minister’s Office, Notice No. 40 of 1995). All efforts were made to minimize animal suffering.

### 2.2 Animals

A total of twelve 6-to 7-week-old male ICR mice (Japan SLC, Japan) were used for recording of neural activity. They were housed in groups under conditions of controlled temperature and humidity (22 ± 1 °C, 55 ± 5%) and maintained on a 12:12-h light/dark cycle (lights off from 8:00 a.m. to 8:00 p.m.) with *ad libitum* access to food and water. Mice pseudo-randomly divided into two groups: the vehicle group *vs*. the theanine group. All mice were weighed at least every three days. The vehicle group was given tap water for daily drinking for three weeks, while the theanine group received tap water containing theanine (see the next section). The amount of water intake was also weighed at least once in three days. Mice were acclimated to experimenters via daily handling at least every three days before experiments.

### 2.3 Drug

L-theanine (C_7_H_14_N_2_O_3_; 326-93463, FUJIFILM Wako, Japan) was dissolved in tap water at a concentration of 40 mg/ml for everyday water intake in the theanine group. Tap water (*i*.*e*., 0 mg/ml theanine) was used as the vehicle control solution.

### 2.4 Surgery

A recording interface assembly was prepared as previously described [26,27,40–42]. In short, the assembly was composed of an electrical interface board (EIB) (EIB-36-PTB, NeuraLynx, USA) and shell and core bodies custom-made by a 3-D printer. The EIB had a sequence of metal holes for connections with wire electrodes. A given individual hole was conductively connected with one end of the insulated wire (∼ 1 cm) using gold pins (for attachment), whereas the opposite end was soldered to a corresponding individual electrode during surgery.

The basic procedure of the stereotaxic surgery was conducted in accordance with our previous literature [25–27,41–43]. General anesthesia was induced and maintained with 4% and 1–2% isoflurane gas, respectively, with careful inspection of the animal’s condition during the whole surgical procedure. Veterinary ointment was applied to the mouse’s eyes to prevent drying.

After complete anesthesia was confirmed, a mouse was laid on its stomach and mounted onto a stereotaxic apparatus (SR-6M-HT, Narishige, Japan) [26]. The scalp was then removed with a surgical knife and scissors. Circular craniotomies with a diameter of approximately 1.0 mm were performed using a high-speed dental drill (SD-102, Narishige). Penetrative nichrome wires (762000, A-M Systems, USA) were stereotaxically implanted unilaterally into the frontal cortex (1.5 mm anterior and 0.5 mm lateral to bregma) and dorsal hippocampus (2.0 mm posterior and 2.0 mm lateral to bregma) to record LFPs. Two stainless-steel screws were additionally implanted into the bone above the cerebellum as ground and reference electrodes. Each of the open edges of ground and reference electrodes was soldered to the corresponding open edge of insulated wires attached to the recording interface assembly. This assembly including all electrodes was secured to the skull using dental cement.

Following surgery, each mouse was allowed to recover from anesthesia and was housed individually with free access to water and food. For the first 3–5 days after surgery, the health condition of animals was carefully checked every day.

### 2.5 Apparatus

The procedure of a novel object recognition task was previously described [13–15]. The behavioral task took place in an open square field measuring 30 cm in width, 30 cm in depth, and 30 cm in height. The walls and floor of the open field were made of translucent corrugated plastic painted black. Each of the four inner walls was independently colored partially with green, red, and yellow tape so that mice could recognize their azimuth in the field. The eight kinds of objects used in this test consisted of [two colors: orange, white] × [four shapes: corn, pyramid, dome, snowman] [14,15]. The base area and height of the objects were approximately 7.1 cm^2^ and 4.0 cm, respectively. A web camera (MCM-303, Gazo, Japan) was placed above the apparatus to monitor the animals’ behavior, and the camera’s strobe signals were transmitted to a data acquisition system (CerePlex Direct, Blackrock Neurotech, USA; see the following sections for details) to synchronize with electrophysiological signals using a custom-made application.

### 2.6 Behavioral tests

Mice in each group were allowed to freely explore the open field (without any objects inside) for 10 min per day and acclimatized to the open field for 4 days. After sufficient acclimatization, the novel object recognition test including training and test sessions took place with a dim light [13,44]. The mouse was tethered to the data acquisition system (CerePlex Direct). In the training session, two identical objects were placed in two of the four quadrants of the open field. A mouse was allowed to freely explore the open field for 5 min and then returned to its home cage. One of the two objects was replaced with another novel object in the same location. Sixty min after the training session, the mouse was allowed to freely explore the field for 5 min. The objects used in each session were predetermined by random selection, and their locations were randomized across mice [13].

Behavioral tests were conducted mostly during the nocturnal period for mice (*i*.*e*., 8 a.m. to 8 p.m.) so that they could behave actively. Every time each behavioral test was finished, the apparatus was wiped, cleaned, and disinfected using 70% ethanol to eliminate as much of the residual mouse scent as possible before the next mouse was placed in the apparatus.

### 2.7 Behavioral electrophysiology

The EIB of the recording interface assembly was connected to a digital headstage (CerePlex μ, Blackrock Neurotech), and the digitized signals were amplified and transferred to the data acquisition device (CerePlex Direct) via interface cables. Electrophysiological signals were digitized at a sampling rate of 2 kHz. LFPs were first recorded for 5 min during the training session, after which mice were returned to the home cage. Sixty hours after the training session, the signals were recorded for 5 min during the test session.

### 2.8 Histology

After the recordings, animals were anesthetized by intraperitoneal injection of an overdose of urethane and transcardially perfused with 0.01 M phosphate-buffered saline (PBS; pH 7.4) and 4% paraformaldehyde (PFA) in 0.01 M PBS, followed by decapitation [16,45–53]. The brains were soaked overnight in 4% PFA for post-fixation and coronally sectioned at a thickness of 100 μm using a vibratome. Serial slices were mounted on glass slides and processed for cresyl violet staining or fluorescence staining [42,43,45–47,50]. For fluorescence staining, the sections were rinsed in 0.01 M PBS containing 0.3% Triton X-100 (called PBST) for 60 min, incubated with 0.1% NeuroTrace™ 435/455 Blue Fluorescent Nissl Stain (N21479, Thermo Fisher Scientific, USA) in PBST for 90 min, and coverslipped with an embedding medium, CC/Mount™ (K 002, Diagnostic BioSystems, USA). Stained images were acquired using a microscope (BZ-X710, Keyence, Japan) [47]. The positions of electrodes were confirmed by identifying tracks in the histological tissue. Data were discarded in the subsequent analysis if the electrode position was outside the target brain region.

### 2.9 Data analysis

All data analyses were performed using custom-made MATLAB (MathWorks, USA). The summarized data are reported as the mean ± the standard error of the mean (SEM). Unless otherwise specified, the null hypothesis was statistically rejected when *P* < 0.05.

The moment-to-moment positions of the mice were tracked by manual detection using ImageJ (National Institutes of Health, USA) [13,42]. For the training session, the time (> 0.4 s) spent around a circular area (∼ 28 cm^2^) around objects that would be unchanged and replaced in the test session was measured and summed to produce T1 and T2, respectively. Similarly, the total time spent around the familiar and novel objects in the test session was defined as T1 and T2, respectively. The discrimination ratio was calculated as (T2 – T1)/(T2 + T1); ideally, the ratio would be zero in the training session. The recording periods dominated by apparent electrical noise caused by grooming, teeth grinding, chewing, and other physical impacts (*e*.*g*., when an animal hit its head on the walls) were manually excluded from the analysis. The LFPs in the hippocampus and frontal cortex were bandpass-filtered at delta (1–4 Hz), theta (6–12 Hz), beta (18–25 Hz), slow gamma (30–50 Hz), and fast gamma (50–80 Hz) bands using fast Fourier transform. For each of the bandpassed LFPs, Hilbert transform was used to calculate the instantaneous amplitude *a*(*t*_*i*_) at every time point *i*; note that the total number of the time points were the product of the total recording time (*i*.*e*., ∼ 300 s for the training and test sessions) and sampling frequency (*i*.*e*., 2,000 Hz) [54]. When each state (*e*.*g*., exploratory behavior for the novel or familiar object in the test session) including *k* time points in total, the average amplitude of the bandpassed LFPs for a given state *S* (*i*.*e*., *A*_state_) was retrieved as follows [14]:

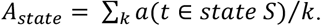

For the training session, the average amplitude was calculated for the exploration of two objects. For the test session, the average amplitude was calculated for each object. A ratio of the average amplitude of given bandpassed LFPs to that of wideband LFPs was multiplied by 100, which was defined as the relative amplitude of the bandpassed LFPs and signified by the unit of %. LFP signals were further convoluted with a complex Morlet wavelet family (bandwidth parameter, 1.5; center frequency, 2) to analyze in a time-frequency domain [26,47,55].

## 3. Results

We chronically fed mice with theanine or tap water (*i*.*e*., vehicle) *ad libitum* for three weeks: two weeks until the surgery day plus one week from the surgery to recording day. For the three weeks of administration, the change in body weight before and after surgery was similar between the vehicle and theanine groups (Fig. 1A), consistent with the almost same amount of water intake between the two groups (Fig. 1B). The body weight on the surgery day was not significantly different between the two groups (40.3 ± 1.1 g (vehicle, *n* = 6 mice) *vs*. 38.9 ± 1.3 g (theanine, *n* = 6 mice), *P* = 0.43, *t*_10_ = 0.82, Student’s *t*-test; Fig. 1A), as was the case with the body weight on the behavioral test day (39.0 ± 0.5 g (vehicle, *n* = 6 mice) *vs*. 36.7 ± 1.6 g (theanine, *n* = 6 mice), *P* = 0.19, *t*_10_ = 1.42, Student’s *t*-test; Fig. 1A). We chronically implanted electrodes into the hippocampus and frontal cortex in mice (see below) and allowed them to perform the novel object recognition task (Fig. 1C, D). The distance traveled by vehicle-treated and theanine-treated mice was not significantly different for the training (27.7 ± 9.2 m (vehicle, *n* = 6 mice) *vs*. 29.2 ± 5.2 m (theanine, *n* = 5 mice), *P* = 0.75, *t*_9_ = −0.32, Student’s *t*-test; Fig. 1E) or test session (23.6 ± 8.5 m (vehicle, *n* = 6 mice) *vs*. 23.6 ± 7.2 m (theanine, *n* = 6 mice), *P* = 0.99, *t*_10_ = 0.01, Student’s *t*-test; Fig. 1E). While the discrimination ratio was not significantly different between the two groups in the training session (−0.16 ± 0.35 (vehicle, *n* = 6 mice) *vs*. 0.16 ± 0.22 (theanine, *n* = 5 mice), *P* = 0.11, *t*_9_ = −1.76, Student’s *t*-test; *P* (vehicle) = 0.31, *t*_5_ = −1.13, one-sample *t*-test *vs*. 0; *P* (theanine) = 0.19, *t*_4_ = 1.60, one-sample *t*-test *vs*. 0; Fig. 1F), the discrimination performance in the theanine group was significantly higher than that in the vehicle group (−0.25 ± 0.26 (vehicle, *n* = 6 mice) *vs*. 0.36 ± 0.23 (theanine, *n* = 6 mice), *P* = 1.50 × 10^−3^, *t*_10_ = −4.33, Student’s *t*-test; *P* (vehicle) = 0.06, *t*_5_ = −2.37, one-sample *t*-test *vs*. 0; *P* (theanine) = 0.01, *t*_5_ = 3.86, one-sample *t*-test *vs*. 0; Fig. 1F), suggesting that chronic intake of theanine enhances memory performance in mice.

**Figure 1.**
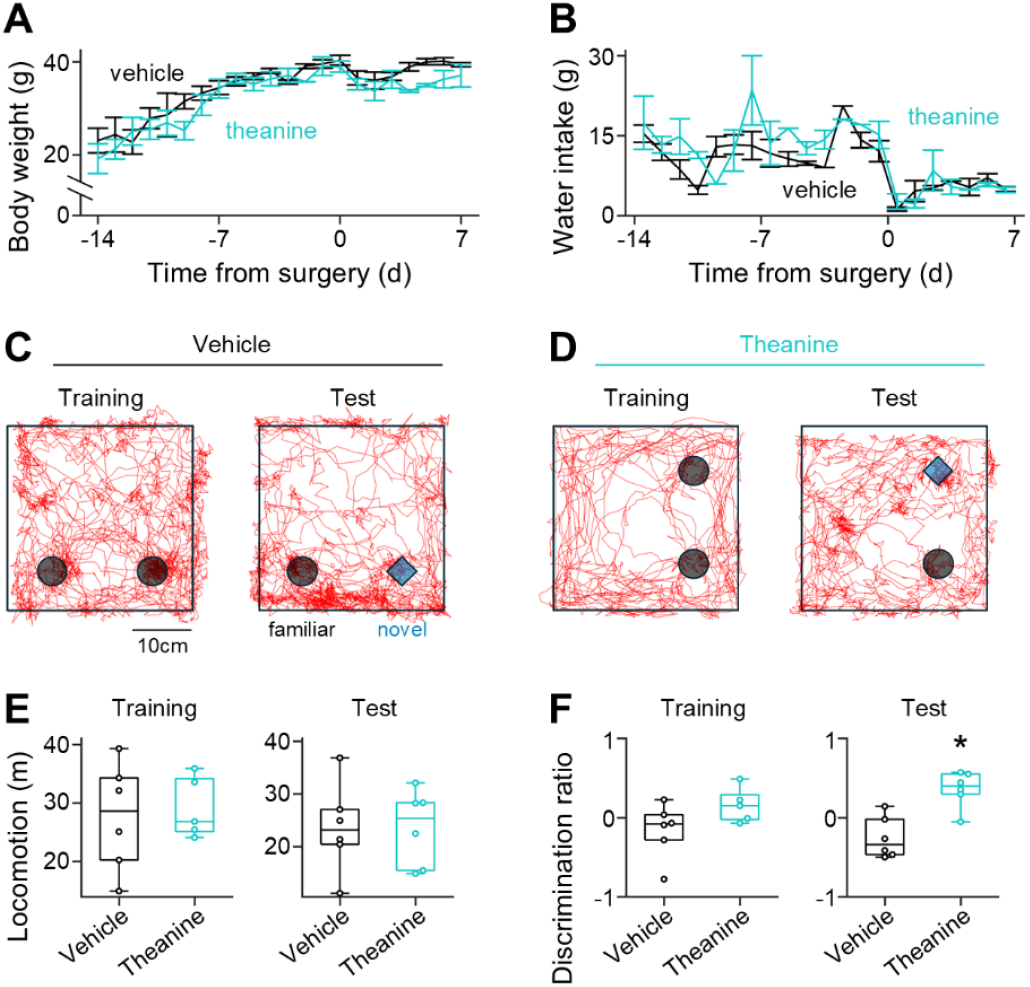
Chronic theanine treatment enhances memory in the novel object recognition task. ***A***, Body weight of vehicle-treated (*black*) and theanine-treated (*turquoise*) mice before and after surgery. ***B***, Same as *A*, but for water intake. ***C***, Representative trajectories (*red*) of a vehicle-treated mouse during the training (*left*) and test (*right*) sessions. Two identical objects (*black*) are located in the training session, but one of them is replaced with another object (*blue*) in the test session. ***D***, Same as *C*, but for a theanine-treated mouse. ***E***, Distance traveled by vehicle-treated (*black*) and theanine-treated (*turquoise*) mice during the training (*left*) and test (*right*) sessions. ***F***, Same as *E*, but for the discrimination ratio.

To seek neural correlates underlying the theanine-increased memory performance, we chronically implanted electrodes into the hippocampus and frontal cortex (Fig. 2).

**Figure 2.**
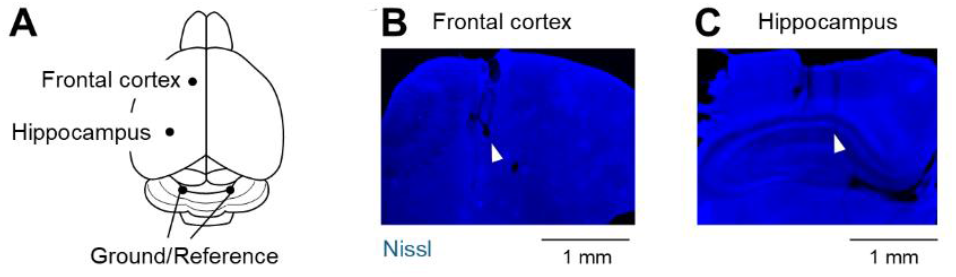
Verification of the location of electrodes chronically implanted into the hippocampus and frontal cortex. ***A***, Top view of the locations of recording and ground/reference electrodes. ***B***, Representative images of Nissl-stained coronal sections including the frontal cortex. Triangles (*black*) denote the tip of the electrodes found in the sections. ***C***, Same as *B*, but for the hippocampus. *Abbreviation*: G/R, ground/reference.

In the training session, we allowed postoperative mice in each group to freely explore two identical objects in an open field and recorded their neural activity from the hippocampus and frontal cortex (Fig. 3A). We evaluated the amplitude of each frequency band of LFPs in the frontal cortex and hippocampus in two groups. The delta, theta beta, low and high gamma amplitude of the frontal LFPs during object exploration was not significantly different between the two groups (delta: 27.1 ± 1.9 % (vehicle, *n* = 6 mice) *vs*. 28.8 ± 2.3 % (theanine, *n* = 5 mice), *P* = 0.60, *t*_9_ = −0.55, Student’s *t*-test; theta: 25.3

**Figure 3.**
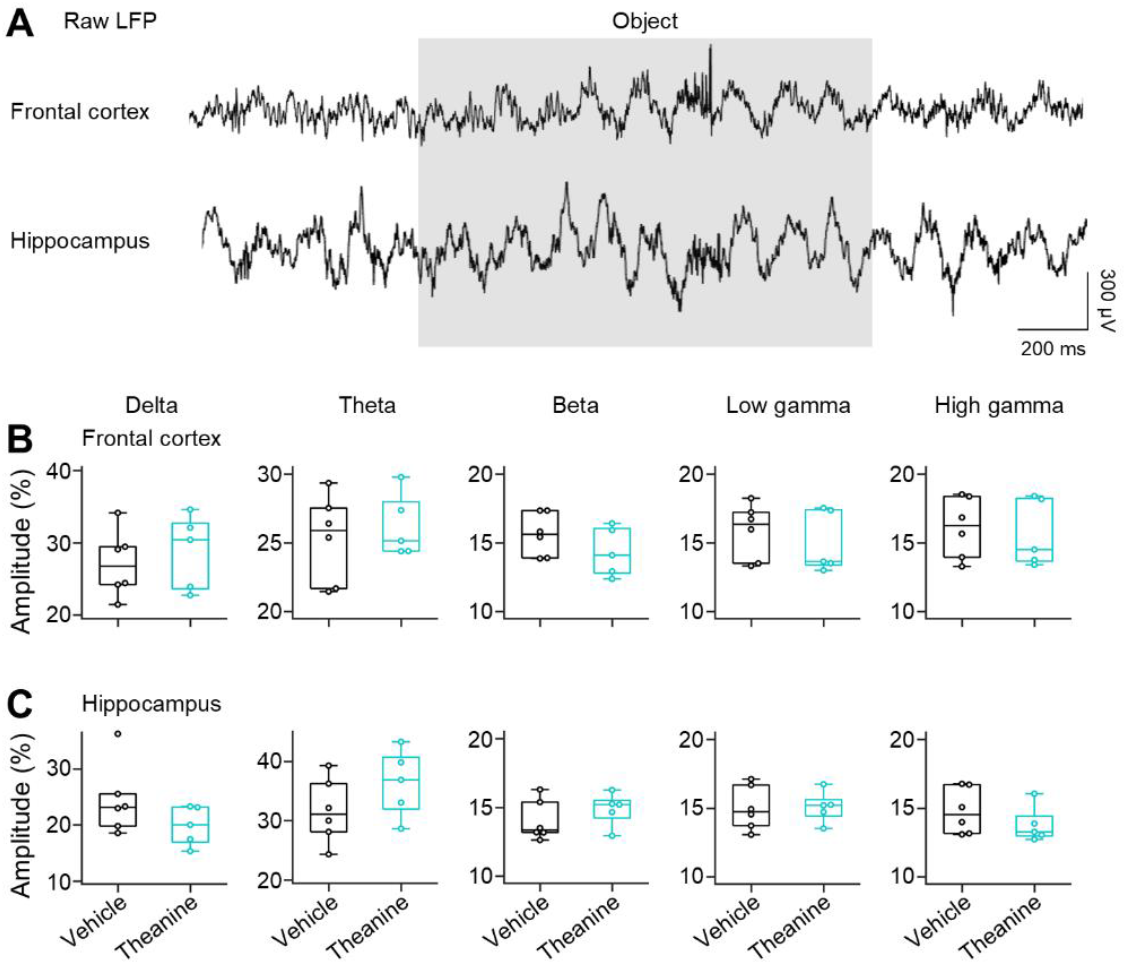
Theanine does not affect any oscillatory components of local field potential signals in the hippocampus or frontal cortex during the training session. ***A***, Representative traces of LFPs in the frontal cortex (*top*) and hippocampus (*bottom*) in a mouse chronically fed with vehicle. The mouse explores one of the objects (*gray*) during the training session. ***B***, Scaled amplitude of delta (*first* (*leftmost*)), theta (*second*), beta (*third*), low gamma (*fourth*), and high gamma (*fifth*) oscillations in the frontal cortex in vehicle-treated (*black*) and theanine-treated (*turquoise*) mice. ***C***, Same as *B*, but in the hippocampus.

± 1.3 % (vehicle, *n* = 6 mice) *vs*. 26.2 ± 1.0 % (theanine, *n* = 5 mice), *P* = 0.61, *t*_9_ = −0.53, Student’s *t*-test; beta: 15.6 ± 0.6 % (vehicle, *n* = 6 mice) *vs*. 14.4 ± 0.8 % (theanine, *n* = 5 mice), *P* = 0.24, *t*_9_ = 1.26, Student’s *t*-test; low gamma: 15.8 ± 0.8 % (vehicle, *n* = 6 mice) *vs*. 15.0 ± 0.1 % (theanine, *n* = 5 mice), *P* = 0.54, *t*_9_ = 0.64, Student’s *t*-test; high gamma: 16.1 ± 0.9 % (vehicle, *n* = 6 mice) *vs*. 15.7 ± 1.1 % (theanine, *n* = 5 mice), *P* = 0.75, *t*_9_ = 0.33, Student’s *t*-test; Fig. 3B), which was comparable with the hippocampal LFPs (delta: 24.4 ± 2.6 % (vehicle, *n* = 6 mice) *vs*. 19.9 ± 1.6 % (theanine, *n* = 5 mice), *P* = 0.19, *t*_9_ = 1.43, Student’s *t*-test; theta: 31.7 ± 2.2 % (vehicle, *n* = 6 mice) *vs*. 36.4 ± 2.6 % (theanine, *n* = 5 mice), *P* = 0.20, *t*_9_ = −1.38, Student’s *t*-test; beta: 14.0 ± 0.6 % (vehicle, *n* = 6 mice) *vs*. 14.9 ± 0.6 % (theanine, *n* = 5 mice), *P* = 0.34, *t*_9_ = −1.01, Student’s *t*-test; low gamma: 15.0 ± 0.7 % (vehicle, *n* = 6 mice) *vs*. 15.1 ± 0.5 % (theanine, *n* = 5 mice), *P* = 0.93, *t*_9_ = −0.09, Student’s *t*-test; high gamma: 14.8 ± 0.7 % (vehicle, *n* = 6 mice) *vs*. 13.8 ± 0.6 % (theanine, *n* = 5 mice), *P* = 0.31, *t*_9_ = 1.08, Student’s *t*-test; Fig. 3C). These results suggest that theanine does not have an impact on the hippocampal and frontal neural activity when the explored objects are the same.

Similarly, we next recorded frontal and hippocampal neural activity in the test session, where one of the objects in the training session was replaced with the novel object; note that the other object was regarded as familiar in the test session (Fig. 4A). Unexpectedly, we found that the theta amplitude in the frontal cortex was significantly enhanced in the theanine group than the vehicle group during the familiar object exploration (theta: 22.1 ± 1.1 % (vehicle, *n* = 6 mice) *vs*. 25.3 ± 0.7 % (theanine, *n* = 6 mice), *P* = 0.04, *t*_10_ = −2.40, Student’s *t*-test; Fig. 4B), while the delta, beta, low and high gamma amplitude in the frontal cortex was not (delta: 32.2 ± 2.8 % (vehicle, *n* = 6 mice) *vs*. 29.5 ± 2.6 % (theanine, *n* = 6 mice), *P* = 0.49, *t*_10_ = 0.72, Student’s *t*-test; beta: 14.5 ± 1.0 % (vehicle, *n* = 6 mice) *vs*. 14.7 ± 0.9 % (theanine, *n* = 6 mice), *P* = 0.89, *t*_10_ = −0.15, Student’s *t*-test; low gamma: 15.5 ± 0.6 % (vehicle, *n* = 6 mice) *vs*. 15.0 ± 1.1 % (theanine, *n* = 6 mice), *P* = 0.75, *t*_10_ = 0.33, Student’s *t*-test; high gamma: 15.7 ± 0.9 % (vehicle, *n* = 6 mice) *vs*. 15.5 ± 1.3 % (theanine, *n* = 6 mice), *P* = 0.89, *t*_10_ = 0.14, Student’s *t*-test; Fig. 4B). Moreover, the beta and low gamma amplitude in the hippocampus was significantly higher in the theanine group than the vehicle group in the familiar object exploration (beta: 13.5 ± 0.6 % (vehicle, *n* = 6 mice) *vs*. 15.7 ± 0.7 % (theanine, *n* = 6 mice), *P* = 0.03, *t*_10_ = −2.47, Student’s *t*-test; low gamma: 15.2 ± 0.5 % (vehicle, *n* = 6 mice) *vs*. 16.6 ± 0.3 % (theanine, *n* = 6 mice), *P* = 0.04, *t*_10_ = −2.35, Student’s *t*-test; Fig. 4C), whereas the other in the hippocampus was not (delta: 28.4 ± 2.4 % (vehicle, *n* = 6 mice) *vs*. 23.3 ± 2.2 % (theanine, *n* = 6 mice), *P* = 0.15, *t*_10_ = 1.58, Student’s *t*-test; theta: 28.4 ± 1.7 % (vehicle, *n* = 6 mice) *vs*. 29.2 ± 2.3 % (theanine, *n* = 6 mice), *P* = 0.78, *t*_10_ = −0.29, Student’s *t*-test; high gamma: 14.5 ± 0.6 % (vehicle, *n* = 6 mice) *vs*. 15.2 ± 0.5 % (theanine, *n* = 6 mice), *P* = 0.35, *t*_10_ = −0.99, Student’s *t*-test; Fig. 4C). In contrast, for novel object exploration, we failed to find any significant differences in the power of either frequency band of frontal LFPs between the two groups (delta: 31.3 ± 3.8 % (vehicle, *n* = 6 mice) *vs*. 28.1 ± 1.2 % (theanine, *n* = 6 mice), *P* = 0.45, *t*_10_ = 0.79, Student’s *t*-test; theta: 22.9 ± 2.2 % (vehicle, *n* = 6 mice) *vs*. 24.6 ± 0.8 % (theanine, *n* = 6 mice), *P* = 0.48, *t*_10_ = −0.74, Student’s *t*-test; beta: 14.6 ± 0.9 % (vehicle, *n* = 6 mice) *vs*. 15.1 ± 2.6 % (theanine, *n* = 6 mice), *P* = 0.62, *t*_10_ = −0.52, Student’s *t*-test; low gamma: 15.6 ± 1.1 % (vehicle, *n* = 6 mice) *vs*. 15.6 ± 0.5 % (theanine, *n* = 6 mice), *P* = 0.94, *t*_10_ = −0.08, Student’s *t*-test; high gamma: 15.7 ± 1.3 % (vehicle, *n* = 6 mice) *vs*. 16.5 ± 0.8 % (theanine, *n* = 6 mice), *P* = 0.60, *t*_10_ = −0.54, Student’s *t*-test; Fig. 4D), as was the case with the hippocampal LFPs (delta: 26.5 ± 3.1 % (vehicle, *n* = 6 mice) *vs*. 21.7 ± 1.7 % (theanine, *n* = 6 mice), *P* = 0.20, *t*_10_ = 1.37, Student’s *t*-test; theta: 29.6 ± 3.1 % (vehicle, *n* = 6 mice) *vs*. 32.4 ± 2.4 % (theanine, *n* = 6 mice), *P* = 0.49, *t*_10_ = −0.72, Student’s *t*-test; beta: 13.5 ± 1.0 % (vehicle, *n* = 6 mice) *vs*. 15.3 ± 0.4 % (theanine, *n* = 6 mice), *P* = 0.14, *t*_10_ = −1.60, Student’s *t*-test; low gamma: 15.7 ± 1.3 % (vehicle, *n* = 6 mice) *vs*. 16.0 ± 0.5 % (theanine, *n* = 6 mice), *P* = 0.80, *t*_10_ = −0.26, Student’s *t*-test; high gamma: 14.7 ± 1.2 % (vehicle, *n* = 6 mice) *vs*. 14.6 ± 0.7 % (theanine, *n* = 6 mice), *P* = 0.94, *t*_10_ = 0.07, Student’s *t*-test; Fig. 4E). These results suggest that theanine-treated mice recognize familiar objects more explicitly than control mice at the level of neural activity, paradoxically leading to improved performance of object discrimination.

**Figure 4.**
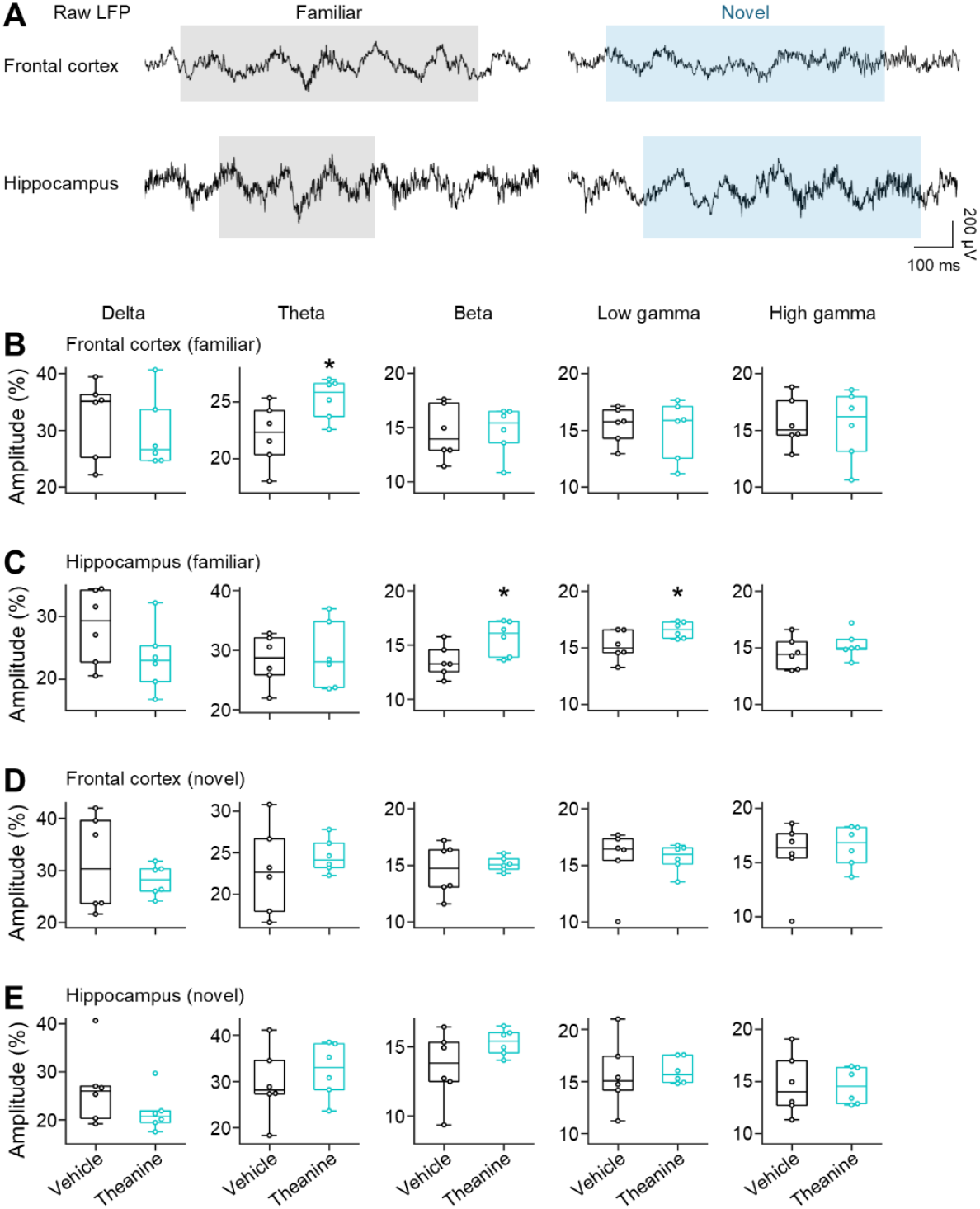
Chronic theanine treatment enhances beta and low gamma oscillations in the hippocampus and theta oscillations in the frontal cortex for object familiarity. ***A***, Representative traces of LFPs in the frontal cortex (*top*) and hippocampus (*bottom*) in a vehicle-treated mouse that explores novel (*blue*) and familiar (*gray*) objects. ***B***, Scaled amplitude of delta (*first* (*leftmost*)), theta (*second*), beta (*third*), low gamma (*fourth*), and high gamma (*fifth*) oscillations in the frontal cortex in vehicle-treated (*black*) and theanine-treated (*turquoise*) mice exploring the familiar object. ***C***, Same as *B*, but for the hippocampus. ***D***, Same as *B*, but for the novel object. ***E***, Same as *D*, but for the hippocampus.

We questioned the temporal order of the enhancement of the frontal theta and hippocampal beta and low gamma oscillations during the familiar object exploration of theanine-treated mice (Fig. 5). We determined the onset of augmentation of each frequency band of LFPs based on the wavelet spectrogram (Fig. 5A–C) and defined the latency as the time from the familiar object exploration to the LFP onset; however, the latencies were not significantly different among the three types of oscillations (0.73 ± 0.14 s (frontal theta, *n* = 18 onsets), 0.66 ± 0.11 s (hippocampal beta, *n* = 22 onsets), 0.61 ± 0.09 s (hippocampal low gamma, *n* = 27 onsets), *P* = 0.72, *F*_2,64_ = 0.33, one-way ANOVA; Fig. 5D). These results indicate that the frontal theta and hippocampal beta and low gamma oscillations are enhanced almost simultaneously.

**Figure 5.**
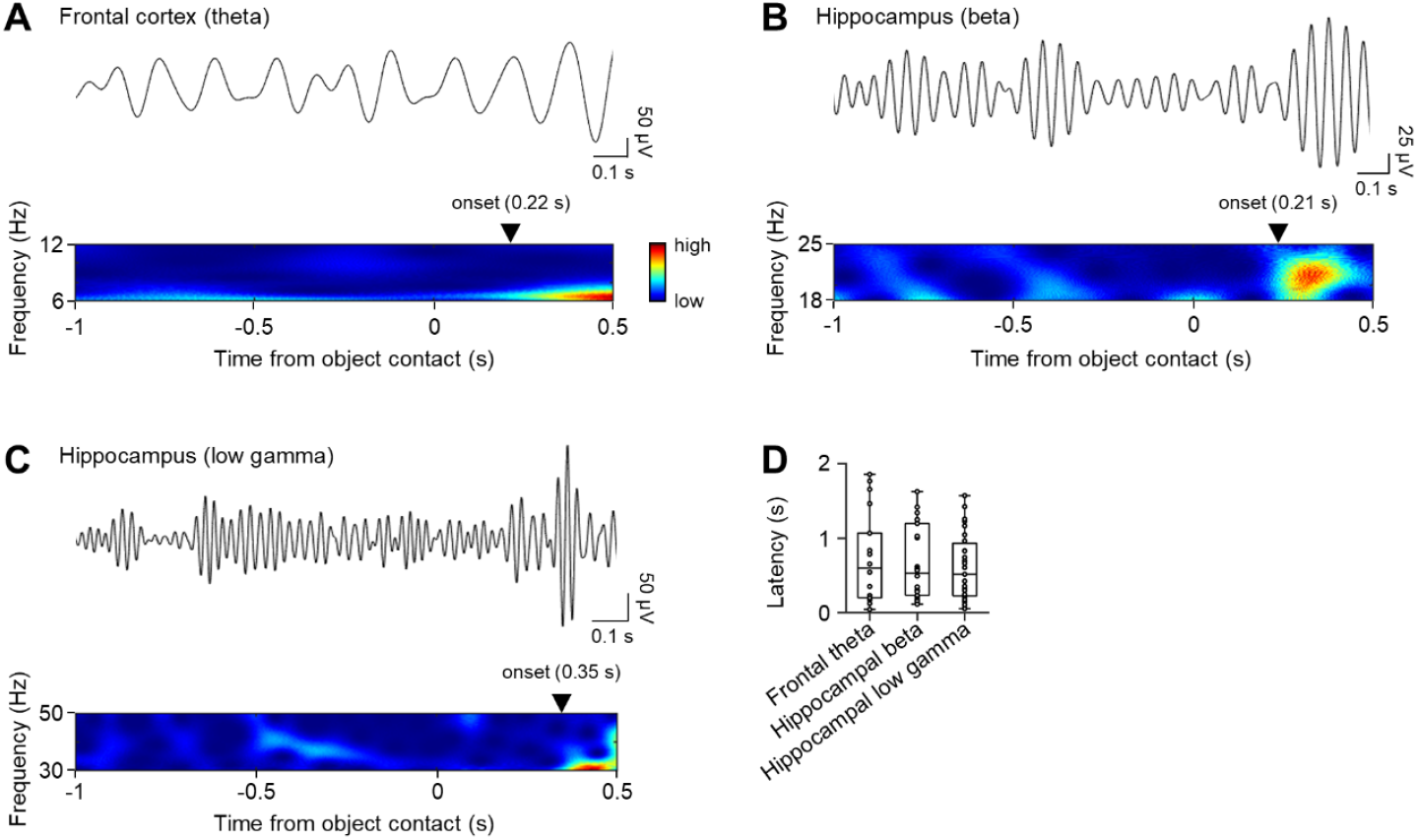
Frontal theta and hippocampal beta/low gamma oscillations in theanine-treated mice for object familiarity are simultaneously enhanced. ***A***, A representative trace (*top*) and wavelet spectrogram (*bottom*) of frontal theta oscillations before and after theanine-treated mice contact familiar objects. The value denotes the latency from the contact with familiar objects to the onset of enhanced oscillations (indicated by the triangle). ***B***, Same as *A*, but for hippocampal beta oscillations. ***C***, Same as *A*, but for hippocampal low gamma oscillations. ***D***, Comparison of latencies of the three bandpassed oscillations.

## 4. Discussion

Here, we found that chronic consumption of theanine enhanced object discrimination in mice. We sought to understand neural activity underlying the enhanced discrimination in theanine-treated mice and recorded LFPs in the hippocampus and frontal cortex during the novel object recognition task. Enhanced memory performance has often been associated with the augmented activity of neurons during the novel object exploration [14,15,54], but unpredictably, theanine boosted the hippocampal beta and low gamma oscillations and frontal theta oscillations for object familiarity. Wavelet analysis confirmed that the three subtypes of oscillations occurred almost simultaneously, raising the possibility that (i) the hippocampus and frontal cortex were regulated by the shared upstream region, or (ii) the two regions were just independently activated.

We found the enhancement of theta oscillations in the frontal cortex in the theanine-treated mice for familiar objects. Extracellular theta oscillations in the frontal cortex have often been considered to be engaged by the interplay with the hippocampus [56,57]. These theta oscillations in the prefrontal cortex and hippocampus are coherent especially when working memory is required [56], and the theta coherence between the two regions is increased by local injection of dopamine [58]. In other words, the frontal theta oscillations are possibly modulated by dopamine. Although it has been poorly understood how dopamine specifically modulates theta oscillations, theanine increases dopamine levels [31], which modulate frontal theta oscillations. On the other hand, we have not found any differences in theta amplitude in the hippocampus between the vehicle-treated and theanine-treated mice. This might be accounted for by the difference in dominance of dopamine D1 and D2 receptors between the hippocampus and frontal cortex in mice [59]. Collectively, when the mice here explore the familiar object in the test session, they may engage their working memory and recall the information of the previously encountered item, which is accompanied by frontal theta oscillations. These frontal theta oscillations are enhanced by theanine, possibly via increased dopamine levels in the brain.

In addition to frontal theta oscillations, we observed enhanced beta and low gamma oscillations in the hippocampus of theanine-treated mice for familiarity. Hippocampal beta oscillations are often observed in mice performing behavioral tasks [18,60,61] and brain slices [62,63]. Mechanisms for generation of beta oscillations have been explored primarily in the rodent hippocampal slices [62]. The plausible model of beta rhythm generation is supported by neural circuits that consist of pyramidal neurons and inhibitory interneurons [64]. According to this model, beta oscillations occur when GABAergic interneurons keep oscillating at gamma frequencies simultaneously with coherent firing of excitatory neurons. The action potentials constantly skip one or two gamma cycles; in other words, firing of excitatory neurons arises during every second or third cycle of gamma rhythms generated by inhibitory neurons [65,66]. The interaction between inhibitory interneurons and excitatory neurons results in net oscillations in gamma and beta frequency bands. In this sense, gamma oscillations are augmented by theanine treatment, because it increases GABA levels in the brain [31,34]. Moreover, hippocampal beta rhythms may be enhanced by activation of hippocampal excitatory neurons. We assume that such activation in the hippocampus is caused by direct or indirect synaptic input of upstream regions including the perirhinal cortex [67], a pivotal region for encoding familiarity [68].

From psychological viewpoints, the object “recognition” in this study may include familiarity and recollection [68]. Generally, a familiarity discrimination system is centered on the perirhinal cortex, whereas a recollective system is based on the hippocampus [69]. According to this psychological theory, augmented extracellular oscillations in the hippocampus for familiar objects observed here are weird; however, chronic theanine administration acts on various receptors, which dynamically modifies neural transmission and circuitry responsible for recognition. Further large-scale physiological recording aimed at multiple regions such as the perirhinal and entorhinal cortices as well as the hippocampus and prefrontal cortex will precisely reveal mechanisms underlying theanine-boosted neural activity for familiarity and deepen our comprehensive understanding of recognition memory.

## Acknowledgments

This work was supported by JST ERATO (JPMJER1801), the Institute for AI and Beyond of the University of Tokyo, JSPS KAKENHI (22K21353), AMED Brain/MINDS 2.0 (24wm0625207s0101; 24wm0625401h0001; 24wm0625502s0501) (to Y. Ikegaya), JSPS KAKENHI (25K18705), and grants from KOSÉ Cosmetology Research Foundation and Hachiro Honjo Ocha Foundation (to N. Matsumoto).

## Author contributions

Y.I. and N.M. conceived the project and designed the study; K.W. performed the experiments; K.W., K.T., T.N. and S.I. analyzed the data; Y.I. and N.M. oversaw and managed the project; K.W., Y.I., and N.M. wrote and sophisticated the manuscript. All approved the final version of the manuscript.

